# Individual Differences in Color-Induced Visual Discomfort Reveal a Pupillary Dissociation Across Stimulus Dimensions

**DOI:** 10.64898/2026.07.17.739245

**Authors:** Ron Y. Meidan, Yoram S. Bonneh

## Abstract

Visual discomfort (VD) is influenced by both spatial structure and chromatic context. Striped patterns are well-established triggers of discomfort and autonomic responses. In previous work, we showed that higher spatial frequencies and larger patterned areas elicit stronger pupillary constriction and greater discomfort, and that individuals with higher overall discomfort show shallower maximum constriction. The present study examined whether similar relationships appear when spatial structure is held constant, measured background luminance is kept within a narrow range, and the chromatic background varies.

Participants viewed black horizontal stripes on 12 near-isoluminant colored backgrounds. The CIE76 color difference (ΔE) ranged from 36 to 112 and was indexed as the CIELAB distance from the black stripe pattern. Pupil size was continuously recorded and discomfort ratings were collected after each trial. Across the colored backgrounds, more uncomfortable stimuli evoked stronger pupil constriction, even though luminance was held nearly constant. As in our spatial-frequency study, this stimulus-level increase did not translate into stronger constriction among observers reporting higher overall discomfort: participants who rated the stimuli as more uncomfortable overall showed shallower constriction. This pattern was captured by the maximum-constriction response, which differentiated high-from low-discomfort observers and was significantly associated with individual discomfort ratings.

Together with our previous findings, the results suggest that pupil responses scale along the tested stimulus axis, whereas individuals reporting greater visual discomfort exhibit less pronounced maximum constriction.

**Highlights:**

- Chromatic background modulated discomfort and pupil responses at similar luminance.
- Across backgrounds, higher discomfort ratings tracked stronger pupil constriction.
- Across observers, higher mean discomfort tracked weaker pupil constriction.
- This two-level dissociation recurs across spatial and chromatic manipulations.
- Pupillometry may complement subjective reports of visual discomfort.

## 1. Introduction

Visual discomfort (VD) is an aversive subjective response to visual stimulation, often elicited by high-contrast, repetitive, or spatially periodic patterns. Striped patterns are among the most established triggers of VD and pattern glare, producing discomfort and perceptual distortions in susceptible observers (Wilkins et al. 1984). Such structures are common in everyday visual environments, including lines of text, fabrics, and architectural patterns (Wilkins et al. 2007). Sensitivity to patterned stimulation varies markedly across people and has been linked to clinical and subclinical traits, including migraine, chronic pain, and other conditions marked by sensory vulnerability (Shepherd, Hine, and Beaumont 2013; Ten Brink, Proulx, and Bultitude 2021), as well as subclinical traits such as schizotypy (Torrens et al. 2024). Pattern glare measures are widely used to characterize this susceptibility (Torrens et al. 2024). Assessment of VD, however, still relies mainly on subjective report. Such reports are essential, but an objective physiological measure would complement them, particularly for observers from whom subjective ratings are difficult to obtain, such as young children or non-verbal individuals, and for developing future screening tools.

Pupillometry is one promising candidate. Pupil size is governed primarily by luminance (Lucas et al. 2003; Spitschan et al. 2014), but it is also modulated by non-luminance properties of the stimulus, including spatial structure, patterned stimulation, and color (Barbur and Thomson 1987; Gao et al. 2020; Hu, Hisakata, and Kaneko 2019; Portengen et al. 2023). Importantly, stimuli that induce discomfort can themselves evoke measurable pupillary responses, including stronger constriction (Ayzenberg, Hickey, and Lourenco 2018; Lin et al. 2015). The pupil is therefore a plausible physiological index of both stimulus strength and individual sensitivity to visual discomfort.

We tested this idea directly in a previous study that manipulated the spatial structure of striped patterns (Meidan and Bonneh 2026). Two relationships emerged. At the stimulus level, higher spatial frequencies and larger patterned areas produced both stronger subjective discomfort and stronger pupillary constriction, when luminance was held constant or even reduced. At the observer level, however, the relationship reversed: participants who reported greater overall discomfort showed shallower maximum constriction. Thus, the same pupillary measure scaled positively with stimulus strength but negatively with individual sensitivity, revealing a dissociation between stimulus-level and observer-level effects.

Whether these relationships are specific to spatial structure, or reflect a more general property of the pupil-discomfort link, can be tested by changing the stimulus dimension while leaving the spatial pattern intact. Color offers such a dimension, and each side of the relationship has independent support in the literature. On the discomfort side, larger chromatic differences between regions increase VD and cortical responses (Haigh et al. 2013; Lindquist, McIntire, and Haigh 2021), and spatial variation in chromaticity predicts discomfort across images (Penacchio et al. 2021); this chromatic discomfort also varies across individuals and with color saturation (Haigh et al. 2026). On the pupil side, chroma and color contrast can modulate constriction even when luminance and lightness are controlled (Portengen et al. 2023; Van Leeuwen et al. 2023). Blue and short-wavelength content is especially relevant because it has been linked to discomfort-related visual disturbance: short-wavelength light increases intraocular straylight and can disturb night vision (Castro-Torres et al. 2024).

These literatures have mostly developed separately. Studies of chromatic discomfort typically record subjective VD or cortical responses without concurrent pupillometry, whereas pupillometric studies of color typically measure brightness or chroma rather than VD. Suzuki et al. (2019) provide a useful pupil-based comparison: using equiluminant colored glare illusions, they found that blue stimuli were perceived as brighter, evoked stronger peak pupil constriction, and showed a participant-level association between brightness adjustments and glare-related pupil constriction. However, their subjective measure was perceived brightness, not visual discomfort. It therefore remains unknown whether chromatic variation in a patterned stimulus produces parallel changes in discomfort and pupil constriction, and whether the observer-level pupil-discomfort dissociation observed for spatial manipulations also appears for color.

Here we held the spatial structure constant, kept measured background luminance within a narrow range, and varied the chromatic background. Participants viewed a fixed pattern of black text-like horizontal stripes on colored backgrounds ordered along a single chromatic axis, indexed by the CIE76 color difference (ΔE) between each background and the black stripe pattern. We worked at low lightness as a practical choice: it let us span a wide range of ΔE while also including the saturated blue of specific interest, and keeping measured luminance within a narrow range reduced the likelihood that large luminance differences alone accounted for the pupil responses (full stimulus details are given in the Methods). Increasing ΔE, chroma, and blue-violet content along the axis were each expected to raise both discomfort and pupillary constriction. This allowed us to test individual sensitivity while keeping the spatial foreground unchanged.

We had two aims. First, at the stimulus level, we asked whether this chromatic axis jointly modulates subjective discomfort and maximum pupillary constriction. Because the chromatic axis combined attributes previously linked to discomfort and pupil constriction, this analysis mainly tested whether the manipulation produced the expected stimulus-level pattern. Second, at the observer level, we asked whether individuals reporting greater overall discomfort again show shallower maximum constriction, as in our spatial study. If the same dissociation recurs under this chromatic manipulation, it would suggest that the observer-level association may extend beyond the specific spatial manipulation used previously.

## 2. Methods

### 2.1 Participants

Forty-three adults participated in the experiment (age range: 18-49 years; 33 males, 10 females). Participants were recruited through routine vision exams. Eligibility criteria included no history of migraine or epilepsy and normal or corrected-to-normal visual acuity. Written informed consent was obtained from all participants in accordance with the Declaration of Helsinki, and the study was approved by the Bar-Ilan University Institutional Review Board.

### 2.2 Eye-tracking and apparatus

The apparatus was identical to that used in our previous work (Meidan and Bonneh 2026). Stimuli were presented on a Yoga 7 14ITL5 laptop with a 14-inch FHD IPS display (1920 × 1080). Screen luminance was set to maximum, and automatic luminance adjustments were disabled. At the nominal viewing distance of 75 cm, the 31 cm × 17 cm display subtended approximately 23.4° × 12.9° of visual angle. Binocular pupil size and gaze position were recorded with a Tobii 4C eye tracker at a sampling rate of 90 Hz. Stimulus presentation and data collection were controlled using PSY, an in-house platform for psychophysical and eye-tracking experiments that has been used previously in our laboratory (Bonneh, Adini, and Polat 2015). To approximate a more naturalistic and potentially portable testing setup, participants were tested without a chinrest, and viewing distance was continuously estimated by the eye tracker. Testing took place in a dimly lit room.

### 2.3 Stimuli

Stimuli consisted of black horizontal stripes composed of random digits rendered in a square-edged monospace font, forming a pattern that mimicked dense lines of text (Figure 1). The global spatial geometry, stripe layout, and digit density were kept constant across all experimental conditions, while the specific sequence of digits within the stripes was randomized on each presentation to avoid pattern-specific effects.

**Figure 1.**
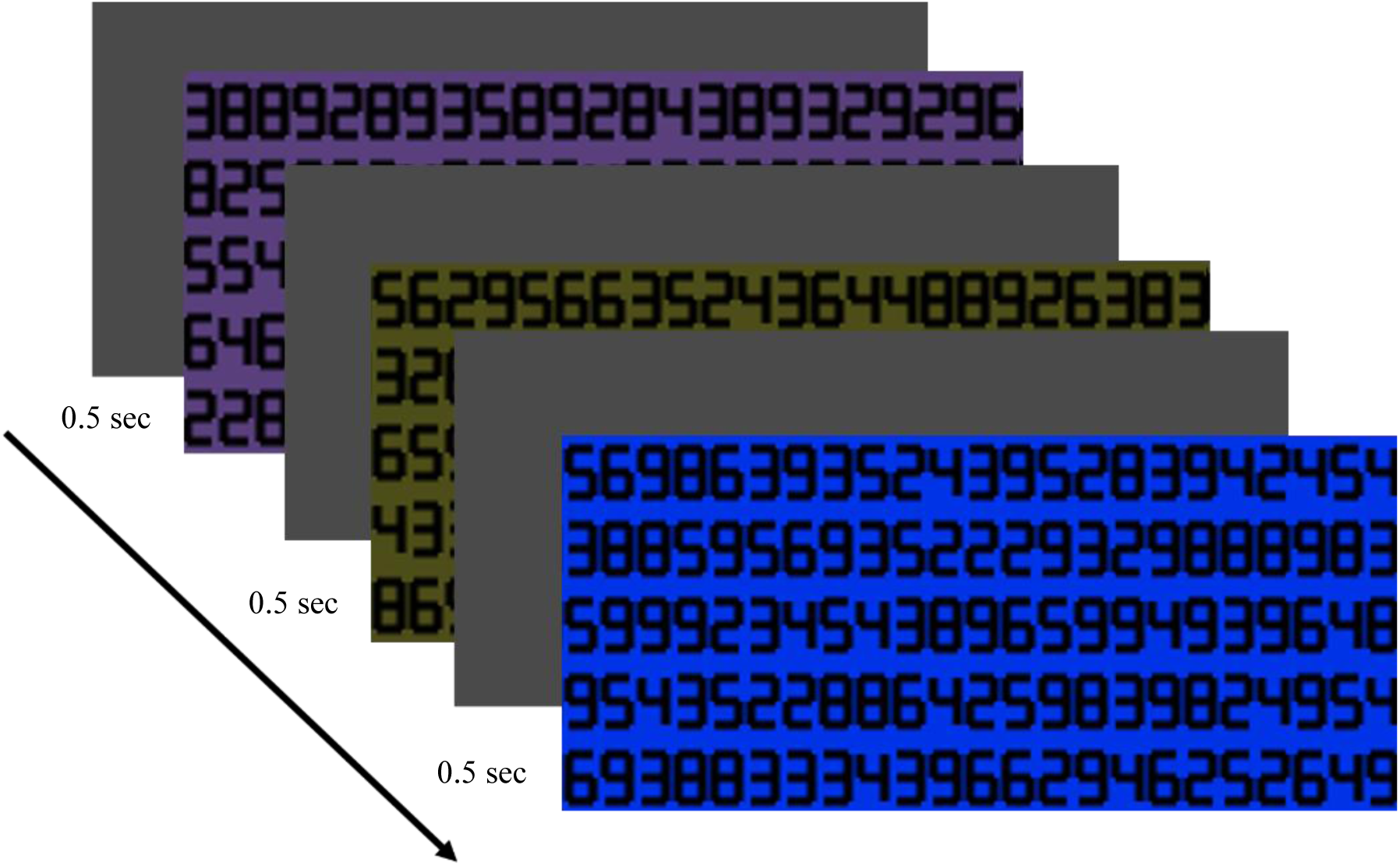
Schematic illustration of the chromatic-background stimulus set. Each background carried the same spatial structure of black digit-like stripes mimicking dense lines of text; the chromatic background varied across conditions. The diagonal arrow indicates the ordering of the twelve backgrounds by ascending CIE76 ΔE from the nominal black stripe pattern. Representative backgrounds from across the ΔE range are shown for illustration. The full set of twelve conditions, with their normalized RGB values, ΔE values, and measured luminance, is summarized in Table 1.

**Table 1.** Background color values, chromatic distance, and measured luminance. Normalized RGB values are expressed on a 0 to 1 scale. ΔE denotes the calculated CIE76 color difference between each background and the nominal black stripe pattern, computed in CIELAB space using a D65 white point. Luminance values indicate the photometric luminance of each background measured with a Minolta LS-100 photometer, showing that background luminance was similar across conditions.

| Stimulus index | R | G | B | $\Delta E$ to black | Luminance (cd/m<sup>2</sup>) |
| --- | --- | --- | --- | --- | --- |
| 1 | 0.20 | 0.30 | 0.40 | 36 | 21.53 |
| 2 | 0.40 | 0.25 | 0.22 | 37 | 21.49 |
| 3 | 0.30 | 0.30 | 0.10 | 44 | 22.03 |
| 4 | 0.39 | 0.25 | 0.09 | 44 | 22.06 |
| 5 | 0.35 | 0.25 | 0.49 | 51 | 20.27 |
| 6 | 0.47 | 0.16 | 0.40 | 55 | 21.18 |
| 7 | 0.05 | 0.34 | 0.00 | 62 | 21.59 |
| 8 | 0.54 | 0.09 | 0.39 | 63 | 22.21 |
| 9 | 0.54 | 0.00 | 0.47 | 73 | 21.51 |
| 10 | 0.50 | 0.00 | 0.60 | 86 | 21.51 |
| 11 | 0.40 | 0.10 | 0.70 | 94 | 20.22 |
| 12 | 0.00 | 0.20 | 0.90 | 112 | 21.30 |

The background colors were selected within the sRGB gamut to span increasing ΔE values from the black stripe pattern while maintaining similar luminance across conditions. Low-ΔE backgrounds included blue-grey, brown, and olive tones. With increasing ΔE, the backgrounds became more chromatically saturated, including magenta and green at intermediate values and purple to blue-purple tones at the highest values. Backgrounds were placed at low lightness (L* ≈ 32) because a saturated blue, which is attainable only at low luminance, could then be included in the set while all backgrounds remained near-isoluminant; this low-lightness range also made it possible to span a wide range of ΔE from the black stripe pattern.

Chromatic distance was quantified as ΔE, defined as the CIE76 Euclidean distance in CIELAB space between each background and the nominal black value of the stripe pattern. Because the pattern elements were black, the reference point was defined as L* = a* = b* = 0. We index the manipulation by ΔE from black rather than by background chroma because ΔE captures the figure-ground contrast between the colored background and the black stripe pattern, whereas chroma describes the background alone. Because lightness was held near L* ≈ 32, ΔE = √(L*² + C*²) reduces to a monotonic function of chroma, and the two indices order the present stimuli almost identically (Pearson r = 0.997, with identical rank order); here, therefore, this is a labeling choice rather than a different manipulation. CIELAB coordinates were computed from the nominal sRGB values of each background using a D65 white point. Based on these computed coordinates, the backgrounds had similar lightness values, with L* ranging from 30.35 to 34.09 (M = 31.6).

Near-isoluminance was then verified photometrically using a Minolta LS-100 photometer. Measured background luminance was similar across the set, ranging from 20.22 to 22.21 cd/m² (M = 21.4 cd/m², SD = 0.6 cd/m²). Screen luminance in the same display setup was also characterized at the standard viewing distance of 75 cm. Fullfield readings yielded 274 cd/m² for white and 0.08 cd/m² for black, indicating a very low black luminance level for the stripe pattern.

Chromatic distance varied substantially across the set, with ΔE ranging from 36 to 112. This range was chosen to provide a broad color sequence for testing whether subjective discomfort and pupillary responses vary together while keeping measured luminance within a narrow range. Stimuli 3 and 4 shared the same ΔE value (44) despite differing in RGB composition. Although the stimuli are shown in Figure 1 and Table 1 in order of increasing ΔE, presentation order was fully randomized for each participant.

### 2.4 Procedure

Participants were seated at a target viewing distance of 75 cm. Because viewing distance was not fixed, it varied across and within participants. The measured viewing distance, averaged across observers, was 78.0 cm (SD = 7.5 cm; range = 61.4 to 92.1 cm). Each of the 12 background colors described above was presented twice in fully randomized order, yielding 24 trials per participant. On each trial, participants first fixated a central marker. The striped stimulus on its colored background was then presented for 0.5 s while pupil size was continuously recorded. No separate practice block was administered. Participants became familiar with the task and rating procedure during the initial trial. Participants were instructed to maintain fixation and to view the stimulus naturally. After stimulus offset, the display transitioned to a grey screen of similar luminance. Participants then rated the discomfort elicited by the just-presented stimulus using a scale from 0 (no discomfort) to 5 (maximum discomfort). After the rating was entered, the next trial began. The full session lasted 1.9 min on average (SD = 0.4 min).

### 2.5 Pupil preprocessing and response measures

Analyses were performed using data from the left eye, selected a priori as in our previous study (Meidan and Bonneh 2026). Pupil data were processed using the same analysis code as described in our lab’s previous work (Kadosh et al. 2024).

Pupil epochs were extracted from 0.5 s before to 3 s after stimulus onset. Within each epoch, pupil size was expressed as percent change from a per-trial pre-stimulus baseline, defined as the mean pupil diameter during the 500 ms preceding stimulus onset. Trials with insufficient usable pupil data in the response window were excluded from pupil analyses. The primary response metric was the maximum pupillary constriction, defined as the most negative percent change in pupil diameter relative to the pre-stimulus baseline within the 0.5-1.5 s post-stimulus window; negative values indicate stronger constriction. For each participant, percent-change values were averaged across eligible trials separately for each background color. Group-level values were then computed from the available participant-level means for each background. The additional requirement of valid pupil data in the 100 ms immediately preceding stimulus onset was applied only to the post-hoc early-session individual-level correlation analysis.

### 2.6 Statistical analysis

All analyses were performed in MATLAB using the same in-house pipeline cited above. We first examined stimulus-level dose-response relationships between chromatic distance from black (CIE76 ΔE) and both visual discomfort (VD) and maximum pupillary constriction. We then examined group-level differences between observers reporting higher versus lower mean discomfort, followed by time-resolved analyses of the pupillary response and individual-level associations between mean constriction and mean VD. For each observer and outcome measure, within-participant outliers were rejected before averaging and model fitting using a ±2 SD criterion. Unless otherwise noted, the same criterion was applied across all analyses reported here.

Stimulus-level effects were assessed using linear mixed-effects analyses with ΔE as a continuous fixed-effect predictor and participant as a random intercept. Individual-level analyses were assessed using Pearson correlations between each observer’s mean constriction and mean VD. For the individual-level scatter analyses, the fitted line was computed using orthogonal (total least squares) regression, which accounts for measurement error in both variables; reported correlation coefficients are Pearson r. For each linear mixed-effects model, we report β, SE, t, degrees of freedom, and p values. Fixed-effect significance was assessed with the residual degrees-of-freedom method, giving t tests on the residual degrees of freedom (the number of observations minus the number of fixed-effect coefficients); the conclusions are unchanged under the more conservative Satterthwaite approximation.

Group comparisons were first examined using the mean VD ≥ 2.5 split, corresponding to the midpoint of the 0-5 discomfort scale and matching the grouping approach used in our previous work. An exploratory cutoff of VD ≥ 2.8 was then examined.

## 3. Results

### 3.1 Pupillary dynamics over time

The experiment used twelve chromatic backgrounds whose CIELAB color difference from the achromatic reference (ΔE) ranged from 36 to 112. Two stimuli shared the same nominal value (ΔE = 44), yielding eleven unique ΔE levels. As shown in Figure 2, increasing ΔE was associated with deeper and more sustained pupil constriction, consistent with a graded pupillary response along the tested chromatic axis. This time-course pattern motivated the analysis of maximum constriction in the 0.5–1.5 s post-stimulus window.

**Figure 2.**
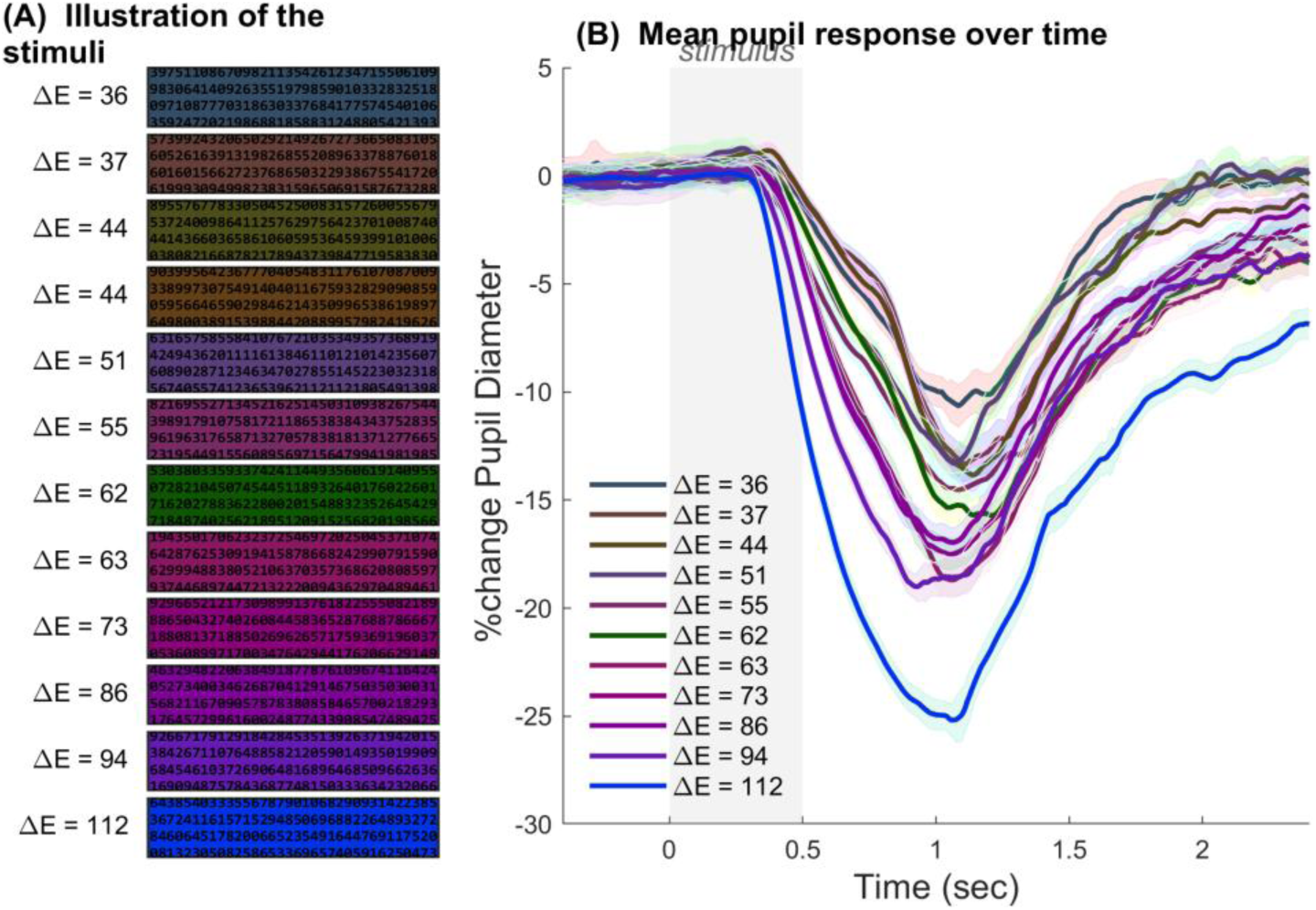
The chromatic-background stimuli and the mean pupillary time course. (A) The twelve background colors used in the experiment, ordered by ascending CIE76 ΔE from the nominal black stripe pattern. Each background displays the same overall spatial structure of black digit stripes; the chromatic background varies between conditions. Stimuli 3 and 4 share the same ΔE value (44). (B) Mean pupillary response over time across 39 to 43 participants per curve, plotted separately for each background. The y-axis shows pupil diameter expressed as percent change from the pre-stimulus baseline (mean over the 500 ms preceding stimulus onset). The grey shaded region from 0 to 0.5 s marks the stimulus-presentation interval. Each curve is color-coded by stimulus ΔE (legend at right); shaded bands indicate ±1 SEM across participants. Higher-ΔE backgrounds produced deeper maximum constriction and a more pronounced sustained response, with the deepest curve corresponding to the highest-ΔE stimulus (ΔE = 112).

### 3.2 Primary dose-response analysis

The dataset comprised 43 observers and 1,020 recorded trials out of 1,032 nominal trials. For one observer, only 12 trials were recorded, resulting in 12 fewer trials than the nominal design. The VD model included 1,020 trials, and the pupil-constriction model included 864 trials with usable pupil data, the reduction reflecting trials lost to blinks or loss of eye-tracker signal in the response window. Both subjective discomfort and pupillary constriction varied systematically along the ΔE-ordered chromatic sequence. In the linear mixed-effects analysis, visual discomfort increased significantly with ΔE (β = 0.016, SE = 0.0015, t(1018) = 10.89, p = 3.3 × 10⁻²⁶). Maximum pupillary constriction became more negative with increasing ΔE (β = −0.15, SE = 0.008, t(862) = −18.70, p = 9.0 × 10⁻⁶⁶). Greater chromatic distance from the black stripe pattern was thus associated with both greater reported discomfort and stronger stimulus-evoked constriction. At the stimulus-mean level, the same monotonic pattern was evident across the eleven ΔE bins (r = 0.840 for visual discomfort and r = −0.939 for constriction).

**Figure 3.**
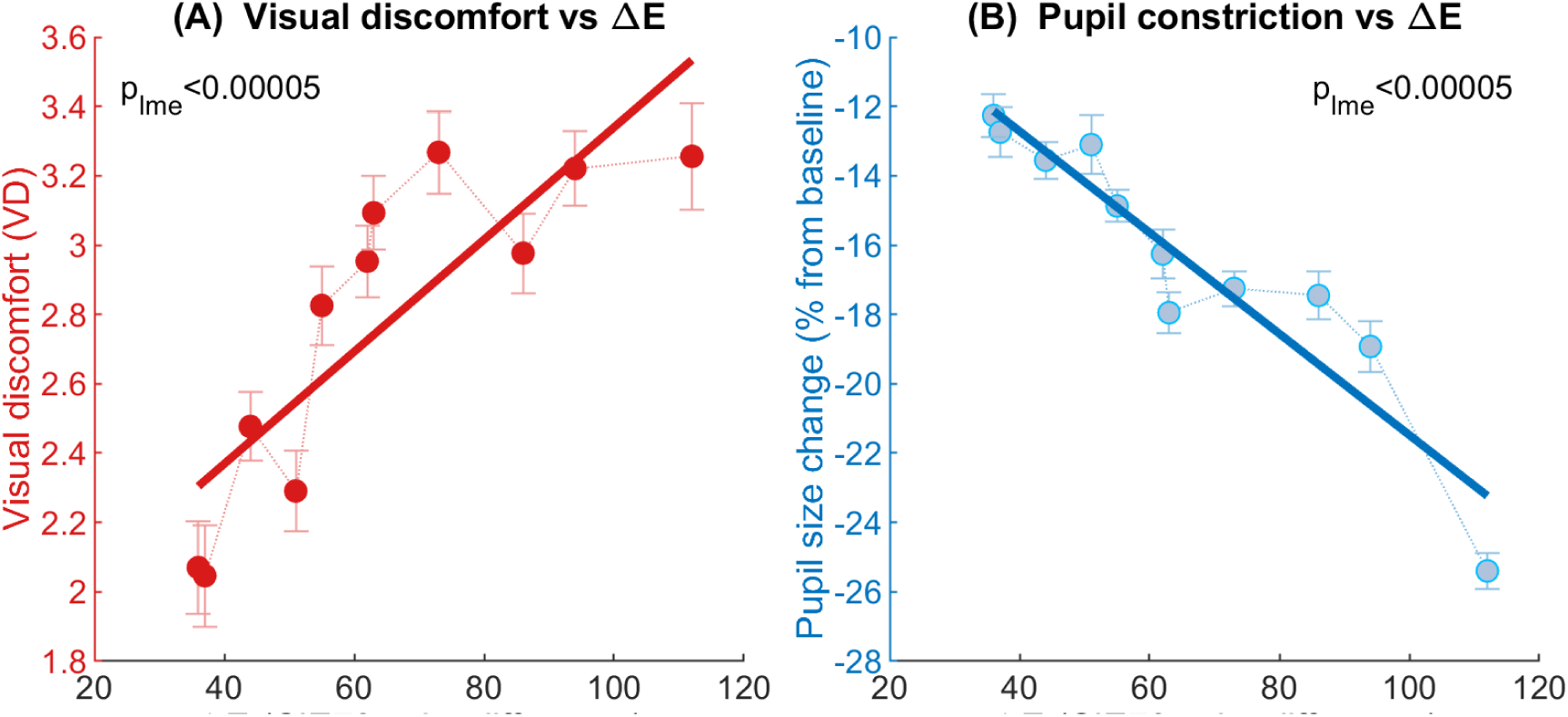
Stimulus-level dose-response in the primary analysis. Group-mean visual discomfort and maximum pupillary constriction plotted as a function of chromatic distance from black (ΔE), with each marker representing the across-participant mean for one of the eleven unique ΔE levels and error bars indicating ±1 SEM. The analysis applies the within-participant ±2 SD outlier-rejection criterion described in the Methods but no additional trial-inclusion filters. The solid line is shown as a descriptive orthogonal (total least squares) regression fit to the plotted means. P-values come from a linear mixed-effects model with ΔE as a continuous fixed effect and participant as a random intercept. (A) Visual discomfort (VD) increased with ΔE: β = 0.016, SE = 0.0015, t(1018) = 10.89, p = 3.3 × 10⁻²⁶; bin means ranged from 2.07 (ΔE = 36) to 3.26 (ΔE = 112). (B) Maximum pupillary constriction, expressed as percent change from the pre-stimulus baseline, became more negative with ΔE: β = -0.15, SE = 0.008, t(862) = -18.70, p = 9.0 × 10⁻⁶⁶; bin means ranged from -12.26% (ΔE = 36) to -25.40% (ΔE = 112).

### 3.3 Group differences in pupillary time course

We first applied the VD ≥ 2.50 split used in our previous study (Meidan and Bonneh 2026), which divided participants into 24 High-VD and 19 Low-VD observers. This split yielded no significant temporal clusters in the pupil time-course analysis; the strongest non-significant cluster occurred from 1100 to 1306 ms post-stimulus (p = .109). We therefore also report an exploratory split using VD ≥ 2.80, which divided participants into a High-VD group (n = 22) and a Low-VD group (n = 21). Because this threshold was chosen after the pre-specified VD ≥ 2.50 split did not reach significance, the group-level time-course result is exploratory and is reported as a robustness check on the continuous individual-level analysis rather than as a confirmatory group difference. A cluster-based permutation test (1,000 permutations) on the full-session pupil time courses identified a single significant cluster spanning 861-2272 ms post-stimulus (p < .001; Figure 4). Throughout this window, the High-VD group showed shallower sustained constriction than the Low-VD group. No significant clusters were present in the pre-stimulus baseline, confirming that the groups were matched before stimulus onset.

**Figure 4.**
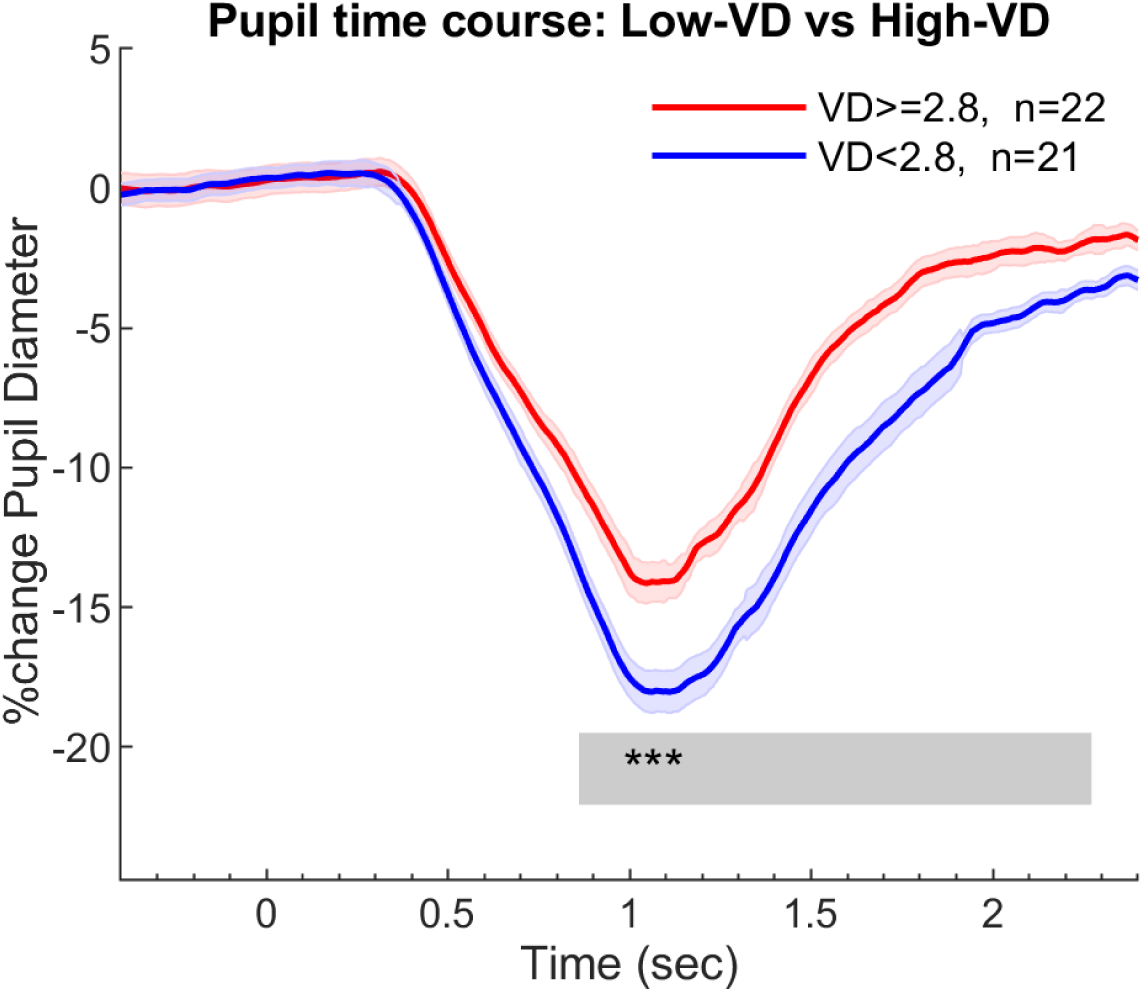
Pupil time course in Low-VD and High-VD observer groups. Mean pupil response over time in observers with lower (Low-VD: VD < 2.80; n = 21) versus higher (High-VD: VD ≥ 2.80; n = 22) mean visual discomfort, expressed as percent change from the pre-stimulus baseline. Shaded bands indicate ±1 SEM across participants. The grey bar at the bottom and the asterisks (***) mark the significant cluster identified by a cluster-based permutation test (1000 iterations), extending from 861 to 2272 ms post-stimulus (p < .001). Throughout this interval, the High-VD group showed shallower sustained constriction than the Low-VD group.

### 3.4 Individual-level association between mean discomfort and mean maximum constriction

At the individual level, mean maximum constriction was positively associated with mean visual discomfort (Figure 5). Because more negative percent-change values indicate stronger constriction, this positive relationship means that observers with higher overall discomfort tended to show less negative mean percent-change values, that is, shallower average maximum constriction. The broad-dataset individual-level correlation was significant (r = +0.44, 95% CI [0.16, 0.65], p = .003, n = 43). This finding is similar to what was observed previously: at the group level, more aversive stimuli produce stronger constriction, but at the individual level, observers reporting higher mean discomfort exhibit shallower average maximum constriction.

**Figure 5.**
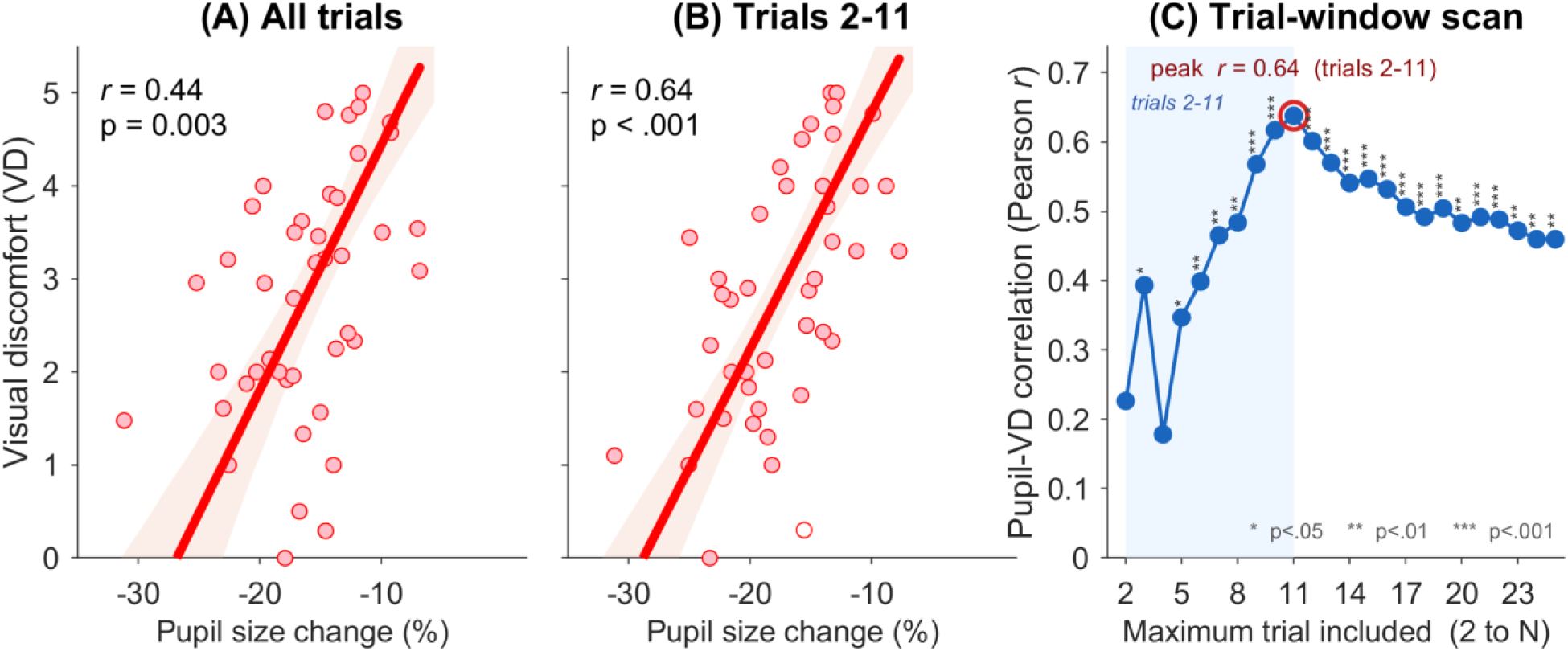
Individual-level pupil-discomfort association. **(A)** All-trials scatter [primary]. Scatterplot of mean visual discomfort (VD) against mean maximum pupillary constriction (expressed as percent change from the pre-stimulus baseline), with one marker per participant (n = 43). No trial-inclusion filter was applied; within-participant outlier rejection at ±2 SD was retained. The red line shows a descriptive orthogonal (total least squares) regression fit; the reported association is the Pearson correlation across observers. r = 0.44, p = 0.003, n = 43. **(B)** Early-session scatter [post-hoc]. Same axes as (A), restricted to trials 2-11, excluding the first trial and requiring a valid pre-stimulus baseline (within-participant ±2 SD outlier rejection). r = 0.64, p = 5.5 × 10⁻⁶, n = 43 (median 9 trials per observer). These filter parameters were identified during exploratory analysis and are therefore treated as a post-hoc sensitivity analysis. **(C)** Trial-window scan. Pearson r between mean VD and mean constriction as a function of the upper trial bound N (cumulative window from trial 2 up to trial N, excluding the first trial and requiring a valid pre-stimulus baseline; ±2 SD outlier rejection throughout). Each marker represents the correlation at one upper bound; significance is indicated by stars (* p < .05, ** p < .01, *** p < .001). The peak (r = 0.64, corresponding to trials 2-11, p = 5.5 × 10⁻⁶) is circled in red.

### 3.5 Exploratory early-session analysis of the individual-level correlation

Higher mean discomfort was consistently associated with shallower mean maximum constriction. The strength of this individual-level association was not fixed in advance but emerged from an exploratory trial-window scan: we computed the correlation over cumulative windows that began at the second trial and extended to successive upper bounds, and the association strengthened from r = 0.44 in the full dataset to a peak of r = 0.64 (95% CI [0.42, 0.79], p = 5.5 × 10⁻⁶) for the window spanning trials 2-11. This window was therefore selected because it maximised the observed correlation, not imposed a priori. The same analysis excluded the first trial, required valid pupil data in the 100 ms immediately preceding stimulus onset, and retained the within-participant ±2 SD outlier-rejection criterion used in the primary analysis. Because the window was chosen to maximise the correlation, this early-session value is treated as a post-hoc, exploratory estimate that requires confirmation in an independent, preregistered study.

### 3.6 Summary of the results pattern

Taken together, the results reveal a coherent pattern across analytic levels. At the stimulus level, increasing ΔE was associated with greater visual discomfort and stronger maximum pupillary constriction. At the observer level, individuals reporting higher mean discomfort showed shallower average maximum constriction, consistent with the individual-difference pattern observed in our previous work. Because the twelve backgrounds were chosen to span an ordered ΔE sequence, the stimulus-level pattern is best read as a graded response along this chromatic axis rather than as an isolated causal effect of ΔE. The individual-level association was significant in the primary all-trials analysis (r = 0.44, p = .003, n = 43) and was stronger in the exploratory early-session window (trials 2-11: r = 0.64, p = 5.5 × 10⁻⁶).

## 4. Discussion

### 4.1 Main findings

The present findings extend our previous spatial work by showing that, under a chromatic manipulation, the relationship between visual discomfort and pupillary constriction also depends on the level of analysis.

### 4.2 Chromatic modulation of discomfort and pupil constriction

Striped and other high-contrast repetitive patterns are among the most established triggers of visual discomfort and pattern glare (Wilkins et al. 1984, 2007; Evans and Stevenson 2008). Their aversiveness has often been discussed in relation to spatial properties, including spatial frequency, contrast, and image statistics that depart from those of natural scenes. In the present study, the same text-like stripe pattern was shown on different chromatic backgrounds while its spatial frequency, stripe density, and patterned area were held constant. Discomfort ratings nevertheless changed systematically with background color. Thus, the discomfort produced by an established spatial trigger was modulated by chromatic context, rather than being determined by spatial structure alone.

This pattern is consistent with broader accounts of visual discomfort as a response to stimulus properties that place unusual demands on visual processing. It is also consistent with previous reports that discomfort increases with chromatic differences between adjacent regions and with chromatic variation in images (Haigh et al. 2013; Lindquist et al. 2021; Penacchio et al. 2021).

The stimulus-level change in discomfort was accompanied by a parallel change in the pupil. Across the same chromatic sequence, maximum constriction deepened in the conditions that produced higher discomfort ratings, measured in the same observers and trials. This is consistent with evidence that the pupil responds to color contrast and chroma when luminance is controlled (Portengen et al. 2023; Van Leeuwen et al. 2023). It also shows that the chromatic modulation of the fixed pattern was expressed both subjectively and physiologically.

This effect should be interpreted in relation to the stimulus set. The backgrounds were ordered by CIE76 color difference from the black pattern, but that ordering also varied chroma and hue direction, with the highest values in the blue-violet range. The stimulus-level result therefore reflects a response to this composite chromatic axis rather than to color difference in isolation. Thus, discomfort ratings and pupil constriction varied together across the chromatic sequence, but the present data do not determine whether this reflects discomfort itself, perceived brightness, or other stimulus-related factors.

### 4.3 Individual differences and the level-of-analysis dissociation

At the observer level, the association reversed: participants who reported greater mean discomfort showed shallower average maximum constriction. Because constriction was expressed as a negative percent change from baseline, this positive association reflects weaker constriction among higher-discomfort observers. The same direction was observed in our previous spatial study (Meidan and Bonneh 2026), arguing against a simple gain account in which discomfort-prone observers merely show amplified pupillary responses to aversive input. Instead, the recurrence of this pattern suggests a distinct observer-level pupil-discomfort relationship, whose possible mechanisms are considered below. This observer-level view complements recent work on individual differences in color-induced visual discomfort (Haigh et al. 2026) by adding a concurrent physiological measure of individual susceptibility.

### 4.4 Relation to perceived brightness

The comparison with perceived brightness helps clarify the interpretation of the pupil response. Suzuki et al. (2019) reported that colored glare illusions, especially blue stimuli, were perceived as brighter and evoked stronger pupillary constriction at equal luminance. At the stimulus level, the present findings are compatible with that result: the higher-ΔE, more blue-violet backgrounds were among the most uncomfortable and produced deeper constriction.

At the observer level, however, the present result differs from what would be expected if discomfort behaved like perceived brightness. In Suzuki et al. (2019), observers who experienced stronger brightness enhancement also showed stronger constriction. Here, observers who reported greater discomfort showed weaker constriction. This difference is not necessarily contradictory, because brightness and discomfort are distinct judgments. Brightness may be more closely tied to perceived light intensity, whereas discomfort may reflect susceptibility to visual stress or the cost of processing an aversive visual pattern.

The present experiment did not include brightness ratings. It therefore cannot determine whether perceived brightness contributed to the stimulus-level pupil response or whether brightness and discomfort make separable contributions.

### 4.5 Possible mechanisms

Several mechanisms could account for the inverse observer-level association. Weaker constriction may contribute to discomfort by allowing greater retinal input during aversive stimulation. Alternatively, shallow constriction and high discomfort may both arise from individual differences in sensory or autonomic regulation. At the cortical level, an elevated excitation-to-inhibition ratio has been proposed as one framework for individual differences in susceptibility to visual discomfort (Penacchio et al. 2023).

Related findings provide context for this interpretation. Reduced constriction amplitudes have been reported in autism spectrum disorder and interpreted in relation to parasympathetic modulation (Daluwatte et al. 2015). Altered pupillary responses to chromatic stimuli have been reported in ADHD (Duque-Chica et al. 2025). In migraine, altered pupillary light responses have been associated with photophobia and disease severity (Cortez et al. 2017). In healthy adults, sensory-processing traits have also been associated with weaker pupillary constriction and differences in the pupillary light-reflex waveform (Kanari and Tsutsumi 2026).

### 4.6 Implications and future work

The findings support pupillometry as an objective complement to subjective reports of visual discomfort. Maximum constriction was sensitive to the chromatic stimulus sequence and was also associated with differences between observers. This dual sensitivity is useful, but it also means that pupil responses must be interpreted according to the level of analysis. Stronger constriction can mark a stronger stimulus-driven response, whereas weaker average constriction may characterize observers who report more discomfort.

Future work should address three questions that the present design could not separate. First, perceived brightness and visual discomfort should be measured together, because the pupil may follow these two subjective dimensions differently. Second, future color sets should test whether the effect of ΔE remains when increasing color difference is not accompanied by increasing blue-violet content. This will likely require testing additional L* ranges, because the low-lightness range used here made it possible to include saturated blue backgrounds but also tied ΔE to chroma and blue-violet content. Third, future studies should ask not only which colors increase discomfort, but also whether some chromatic backgrounds improve comfort relative to an achromatic or black-white baseline, and whether such improvement is reflected in the pupil response.

## 5. Limitations

First, the design did not separate chromatic distance from chromatic direction. Within the low-lightness range used here, color difference from the black pattern was closely tied to background chroma, and hue direction and blue-violet content changed along the same sequence. Across the twelve backgrounds, color difference also covaried with the a* red-green axis and, to a lesser degree, with L*, despite the narrow measured luminance range. The stimulus-level relationship therefore reflects a bundled chromatic manipulation rather than variation along a single color axis. The present data cannot identify color difference, chroma, hue direction, or short-wavelength content as the independent operative factor.

Second, the backgrounds were specified using nominal sRGB values, from which CIELAB coordinates were computed, and their luminance was verified photometrically. Spectral power distributions were not measured. The study therefore cannot separate short-wavelength or melanopsin-related contributions from the other chromatic differences in the sequence. Spectrally controlled stimuli would be needed to test those contributions directly, for example by varying short-wavelength or melanopsin-related stimulation while holding the other color properties as constant as possible.

Third, the observer-level association was present in the broad all-trials analysis and became stronger in the early-session analysis. The present data cannot determine whether this time course reflects adaptation, habituation, task familiarization, fatigue, cumulative discomfort, or another process that changes across trials. The early-session estimate should therefore be confirmed in an independent, preregistered study.

Fourth, observer sensitivity was defined only from discomfort ratings collected within this experiment. Participants did not complete an independent measure of visual sensitivity, such as a pattern-glare inventory, migraine questionnaire, or sensory-processing measure. Terms such as higher-discomfort and lower-discomfort therefore refer only to mean ratings within this study and should not be interpreted as clinical or trait-level classifications.

Fifth, the study used a naturalistic viewing setup without a chinrest, consistent with the goal of approximating a portable testing context. Viewing distance was monitored by the eye tracker, but small changes in head position, viewing distance, or accommodation may have added noise, particularly to the individual-level analyses.

Sixth, the sample was predominantly male. The study was not designed or powered to test sex or gender differences, and this imbalance may limit generalizability. Future studies should recruit a more balanced sample and examine whether the pupil-discomfort relationship differs by sex or gender.

## 6. Conclusion

Changing the chromatic background of a fixed striped pattern systematically modulated both discomfort ratings and maximum pupillary constriction. Across stimuli, higher discomfort was accompanied by stronger constriction. Across observers, however, higher mean discomfort was associated with shallower constriction.

This same two-level structure was previously observed with spatial manipulations, suggesting that stimulus-driven and observer-driven pupil effects are distinct components of the pupil-discomfort relationship. The findings support maximum pupillary constriction as a physiological measure that may complement subjective discomfort ratings, while also showing that the measure cannot be interpreted in a single direction across all levels of analysis.

Future work should determine which chromatic variables drive the stimulus-level effect, whether perceived brightness and discomfort make separable contributions, and whether the observer-level association generalizes to independent measures of visual sensitivity.

## Data Availability

De-identified data supporting the findings of this study are available from the corresponding author on reasonable request for non-commercial research. The data are not deposited in a public repository because the Institutional Review Board approval and participant informed consent did not cover public release of individual eye-tracking records.

## Code Availability

The custom MATLAB analysis scripts and experiment-control code that support the findings of this study are available from the corresponding author upon reasonable request.

## Author Contributions (CRediT)

R.M.: conceptualization, methodology, investigation, data curation, writing - original draft. Y.S.B.: conceptualization, methodology, supervision, writing - review & editing.

## Declaration of generative AI and AI-assisted technologies in the manuscript preparation process

During the preparation of this work, the authors used ChatGPT and Claude to assist with manuscript organization, language editing, clarity, and formatting. After using these tools, the authors reviewed and edited the content as needed and take full responsibility for the content of the article.

